# The Pathological Role and Therapeutic potential of ALDH2 in acrolein detoxification Following Spinal Cord Injury in Mice

**DOI:** 10.1101/2025.06.08.658505

**Authors:** Siyuan Sun, Seth Herr, Anna Alford, Rachel Stingle, Zaiyang Zhang, Timothy Dong, Riyi Shi

## Abstract

Oxidative stress and lipid peroxidation-derived aldehydes, such as acrolein, play a central role in the pathology of spinal cord injury (SCI) and have emerged as promising therapeutic targets. Mitochondrial aldehyde dehydrogenase-2 (ALDH2) is a key oxidoreductase responsible for detoxifying reactive aldehydes. Pharmacological activation of ALDH2 using Alda-1, a selective agonist, has been shown to reduce aldehyde accumulation, alleviate inflammation, and enhance functional recovery in experimental SCI models. However, approximately 8% of the global population carries the ALDH2*2 mutation, which severely impairs this detoxification pathway. In this study, we used a transgenic ALDH2*2 mouse model to investigate the role of ALDH2 in SCI pathology. This model mimics the human ALDH2*2 condition, allowing us to examine the impact of impaired aldehyde clearance on acrolein accumulation and its pathological consequences. We modulated endogenous aldehyde detoxification through both genetic deficiency and pharmacological activation with Alda-1. Our results showed that ALDH2 deficiency led to significantly elevated acrolein levels, which were associated with increased microglial activation, cytokine storm, neuronal loss, demyelination, and tissue damage compared to wild-type (WT) mice. Treatment with Alda-1 enhanced ALDH2 activity and significantly reduced acrolein levels in both ALDH2*2 and WT mice from 2 to 28 days post-SCI. This was accompanied by reduced inflammation, improved preservation of myelin, and marked improvements in locomotor and sensory function, especially in ALDH2*2 mice. Notably, even beyond the traditionally ideal treatment window, Alda-1 treatment remained effective in promoting recovery, particularly in motor function and to a greater extent in ALDH2*2 mice. Our study comprehensively evaluated ALDH2’s role in SCI by both genetically impairing and pharmacologically enhancing its activity, highlighting ALDH2 as a critical modulator of acrolein-mediated damage and suggesting its potential as a therapeutic target, especially for individuals with the ALDH2*2 mutation.

## Introduction

Spinal cord injury (SCI) affects approximately 27 million people worldwide, leading to severe motor, sensory, and autonomic dysfunction that significantly impairs daily activities and quality of life^1,2^. Beyond the initial mechanical trauma, the secondary injury cascade plays a crucial role in exacerbating tissue damage and functional deficits^3,4^. Among these secondary mechanisms, oxidative stress has been extensively implicated as a key driver of neurodegeneration following SCI^5–7^. Excessive production of reactive oxygen species (ROS) induces lipid peroxidation, targeting polyunsaturated fatty acids (PUFAs) within cellular membranes and generating highly reactive aldehydes such as acrolein, 4-hydroxynonenal (4-HNE), and malondialdehyde (MDA)^8–10^. Of these, acrolein is particularly toxic due to its high reactivity, prolonged half-life, and ability to form adducts with proteins, lipids, and DNA, further amplifying cellular damage^11–13^.

Previous studies from our lab have demonstrated the therapeutic potential of anti-acrolein treatments in central nervous system (CNS) trauma and diseases, including SCI, multiple sclerosis (MS), and Parkinson’s disease (PD)^14–17^. Using repurposed acrolein scavengers such as hydralazine, phenelzine analog (PhzA), and dimercaprol, these studies have consistently shown improved outcomes, highlighting the efficacy of neuroprotection through exogenous acrolein scavenging^18–23^. However, despite their benefits, many of these scavengers are associated with undesirable side effects, as they were originally FDA-approved for other conditions^24,25^. This limitation underscores the need to explore alternative therapeutic agents with minimal side effects before advancing to clinical applications.

Mitochondrial aldehyde dehydrogenase-2 (ALDH2) is a crucial oxidoreductase that serves as a primary defense against toxic aldehydes in both humans and rodents^26–28^. Recent studies have highlighted its direct involvement in SCI, demonstrating that pharmacological activation of ALDH2 with Alda-1, a selective ALDH2 agonist, effectively reduces aldehyde accumulation, alleviates inflammation, and enhances functional recovery in experimental SCI models^29,30^. These findings illustrate the essential role of ALDH2 in acrolein detoxification, suggesting that its modulation may be a promising therapeutic strategy for mitigating SCI-induced pathology.

Individuals carrying the ALDH2*2 mutation, which severely impairs aldehyde detoxification, face a significantly higher risk of developing certain cancers, cardiovascular diseases, and neurodegenerative disorders such as Alzheimer’s and Parkinson’s disease— conditions strongly linked to aldehyde accumulation^31–36^. The ALDH2*2 missense variant, best known for causing alcohol flushing reactions, is one of the most extensively studied genetic polymorphisms in humans and is predominantly found in East Asian populations, affecting nearly 8% of the global population^37,38^. Given the growing evidence associating ALDH2*2 with multiple diseases, there has been increasing interest in targeting ALDH2 activation for therapeutic intervention. Studies in cellular and animal models have demonstrated that enhancing ALDH2 activity can mitigate aldehyde toxicity and improve disease outcomes^26,39–42^. However, despite these advancements, the impact of ALDH2*2 on SCI pathology remains under-explored, highlighting a crucial gap in understanding its role in neurotrauma and potential therapeutic strategies.

As such, this study aimed to elucidate the role of ALDH2 and acrolein in SCI pathology by examining a comprehensive range of pathological hallmarks, including inflammation, demyelination, and neuronal loss, as well as the timing of treatment. Using a transgenic murine model (ALDH2*2) that closely mimics the human ALDH2*2 condition, we modulated endogenous acrolein detoxification through genetic deficiency and pharmacological activation with Alda-1. We provided evidence that insufficient ALDH2 activity contributes to an exacerbated pathology compared to mice with a wild-type (WT) background. We have also offered further insights into the mechanisms of reactive aldehydes and the efficacy of acrolein-scavenging treatments for SCI. Interestingly, while boosting endogenous acrolein scavenging brought neurological improvements in both the ALDH2*2 and WT mice after SCI, it was shown to be more effective in individuals with an ALDH2*2 background.

## Materials and Methods

### Animal Experiment

ALDH2*2 human E487K mutant knock-in with a C57/BL6 background mouse line was received as a gift from Dr. Che-Hong Chen at Stanford University. Heterozygous mice were backcrossed to achieve homogeneous ALDH2*2 and WT ALDH2 genetic backgrounds, confirmed by genotyping. The animals were housed in a facility with temperature and illumination control, with free access to food and water ad libitum. Up to five animals were caged together. The animals were acclimated for at least 1 week before surgery. Young adult male ALDH2*2 and wild-type C57BL/6J mice, age matched 10–12 weeks, were randomly divided into the experimental groups: Sham (T9-T10 laminectomy only), SCI, and Alda-1 treatment (SCI with Alda-1 treatment). All experimental procedures mentioned in this paper were approved by the Institutional Animal Care and Use Committee at Purdue University (IUCAC) in October 2023 (approval No. 1909001957). The experimental design and timeline are shown in Fig. 1.

**Figure 1.**
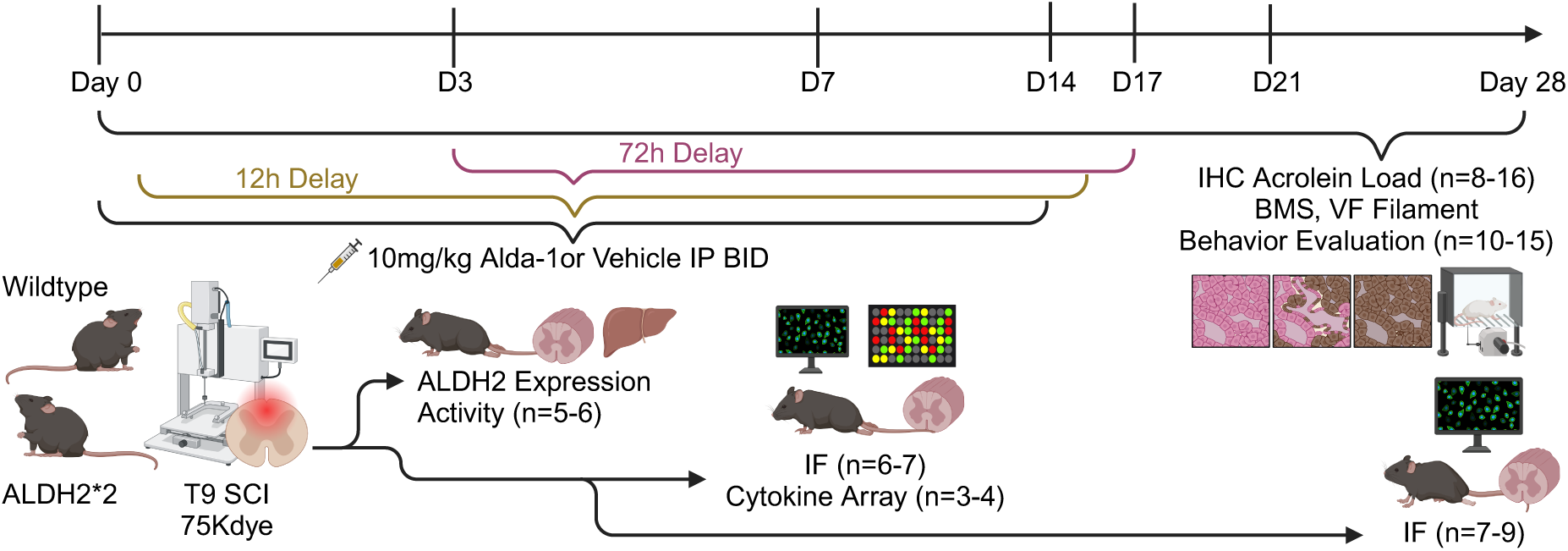
Overview of the experimental timeline. Wild-type and homozygous transgenic ALDH2*2 mice were assigned to sham, spinal cord injury (SCI), and SCI with Alda-1 treatment groups, with interventions beginning on day 0 (six groups in total). In the treatment groups, Alda-1 (10 mg/kg) was administered twice daily for the first two weeks. Behavioral assessments, including the Basso Mouse Scale (BMS) for locomotion and von Frey (VF) filament testing for nociceptive sensitivity, were conducted on day 0 and weekly thereafter until day 28. Additionally, a separate cohort of wild-type and ALDH2*2 mice was used for a delayed treatment experiment (four groups in total), in which the first Alda-1 dose was administered either 12 hours (gold line) or 72 hours (purple line) post-injury, followed by a 14-day treatment regimen.

### Mice Spinal Cord Injury Model

All surgeries were conducted under standard aseptic conditions. Mice were anesthetized via intraperitoneal injection of a ketamine (100 mg/kg) and xylazine (10 mg/kg) cocktail. Anesthesia was confirmed by the absence of a withdrawal response to a rear foot pinch. After shaving and sanitizing the dorsal skin, a longitudinal incision was made to expose the vertebrae at the T9-T10 level. A laminectomy was performed to access the spinal cord at this site. A moderate contusion injury was induced using the Infinite Horizon Impactor (IH impactor) with an impact force of 75 Kdyne^43,44^. Criteria for a successful establishment of the model were as follows: hemorrhage and edema in the injured area; the dura mater remained intact and turned purple; and acute flaccid paralysis (Basso Mouse Scale (BMS) locomotor rating scale of 0-1). Following surgery, the animals were placed on a heating pad for recovery and administered NSAID Carprofen (10 mg/kg subcutaneously) once daily for three days. Bladders were manually expressed twice daily until reflexive control returned. For the sham group, only the laminectomy was performed at T9-T10. Animals that met exclusion criteria were excluded from the study and euthanized immediately following the study protocol.

### Alda-1 Treatment

Alda-1 [N-(1,3-benzodioxol-5-ylmethyl)-2,6-dichlorobenzamide] (Cat# 21555) was purchased from Cayman Chemical Company (Ann Arbor, MI, USA). It was dissolved in 50% dimethyl sulfoxide (DMSO) and 50% polyethylene glycol 400 (PEG-400) with the concentration of 10 mg/ml and administered through intraperitoneal (IP) injection slowly at 30 mins after SCI with corresponding dosages (10mg/kg) and twice a day up to 14 days. Mice in the Sham and SCI groups received an equivalent volume of the vehicle injection. Animals in the delayed treatment groups initially received vehicle injections, followed by the first dose of Alda-1 administered either 12 hours or 72 hours post-injury. Subsequent Alda-1 injections were given twice daily for an additional 14 consecutive days.

### Enzyme Activity Assay

At Day 3 post-SCI, the mice were anesthetized and perfused intracardially with a cold, phosphate-buffered saline (PBS) solution. The fresh spinal cord and liver samples were collected right after to evaluate the ALDH2 enzyme activity levels using a commercial colorimetric kit (#ab115348; Abcam) according to the manufacturer’s protocol and a previous publication. Briefly, tissue lysates were homogenized and centrifuged at 14,000 RPM for 20 minutes at 4°C and diluted to 25 mg/ml (liver) and 50mg/ml (spinal cord). After a series of incubations, the enzymatic activity was measured at 25°C. The nicotinamide adenine dinucleotide hydrogen (NADH) production level was determined spectrophotometrically by monitoring the alterations in absorbance intensity at 450 nm every 30 seconds for 60 minutes using the kinetic mode on a SPECTRAmax plate reader (Molecular Devices, Sunnyvale, CA). The slope of the reaction speed for each group at 5 min was used to calculate relative activity.

### Western Blot

Spinal cord tissues harvested at Day 3 post-injury were stored at –80℃ and used for Western Blot. Briefly, half an inch of the spinal cord tissue from the injury epicenter was homogenized in a 1xRIPA buffer supplemented with a protease inhibitor cocktail, then centrifuged at 14,000 RPM for 30 minutes. Protein concentrations in the supernatant were determined using the Pierce™ BCA Protein Assay Kit (Rockford, IL, USA) and a SPECTRAmax plate reader. Thirty micrograms of protein were mixed with 20% SDS, β-mercaptoethanol (BME), and Laemmli buffer, then loaded onto 15% Tris-HCl gels for electrophoresis. An identical set of samples was loaded onto another gel and run simultaneously, then visualized with Coomassie blue stain (ThermoScientific, 20278) for normalization. The separated proteins were transferred to a nitrocellulose membrane using the Power Blotter-Semi-dry Transfer System (Thermo Fisher Scientific Inc., MA, USA). The membrane was blocked with a 1x casein solution (Vector, #SP-5020), then incubated overnight at 4℃ with primary antibodies, 1:1000 rabbit-ALDH2 (ThermoFisher, PA5-11483). The membranes were subsequently incubated with the 1:2000 secondary antibodies (Vector, #BA-1000) at room temperature for 45 minutes, then treated with VECTASTAIN ABC-AmP reagent (Vector Laboratories, CA, USA) for 10 minutes to amplify the signal. DuoLuX substrate (Vector, #SK-6605) was applied for chemiluminescent detection using the Azure C300 Western blot imaging system (Azure Biosystems, Dublin, CA). Scanning of the Coomassie blue-stained gel and membrane whole protein bands was performed using Azure Sapphire Biomolecular Imager (Azure Biosystems, Dublin, CA). Band intensities were quantified using ImageJ software (NIH, USA).

### Cytokine Expression Assay

Proteome Profiler Mouse Cytokine Array Kit (R&D Systems, ARY006) was used to determine cytokine expression in spinal cord homogenates following the manufacturer’s instructions. Briefly, the mice were anesthetized and perfused intracardially with a cold PBS solution, and spinal cord tissue was harvested at Day 7 post-SCI. Tissues were homogenized immediately, and protein concentrations were measured using the same BCA assay kit as described in the Western Blot section. 450 μg of protein lysate was used for each membrane. Membranes were blocked, incubated, and washed according to standard protocol. Six membranes were exposed to the Azure C300 Western blot imaging system for 10 minutes at the same time. The intensity (pixel density) of each spot on the membrane was quantified using Image J software (NIH, USA), corrected for background intensity, and normalized to the sham group.

### DAB Immunohistochemistry

At Days 2, 7, and 28 post-SCI, spinal cords were extracted after perfusion, and a half-inch segment centering on the injury site was dissected. Samples were stored in 4% PFA for 24 hours, followed by 30% sucrose/PBS solution for 72 hours. Tissue samples were frozen in OTC using a dry ice/isopentane slurry, then cryo-sectioned (Thermo HM525NX) into 25 μm-thick slices. Slices were stored in PBS containing 0.1% sodium azide in 4℃ before biochemical analysis.

The DAB staining and analysis protocol has been described in our previous publications^29,45^. Briefly, spinal sections 1.25 millimeters caudal to the injury epicenter were washed in PBS solution, then placed in 3% H_2_O_2_/water to quench endogenous peroxidase activity. After washing three times in PBS, sections were transferred into 10% normal goat serum for blocking for 1 hour at room temperature, then incubated overnight with an anti-acrolein-lysine antibody (StressMarq; SMC-504D) in 4℃. This was followed by an incubation in biotinylated secondary antibodies (Vector Laboratories) and an ABC avidin/biotin complex solution (Thermo ScientificTM; 32020) before being developed using the DAB Peroxidase (HRP) Substrate Kit, 3,3′-diaminobenzidine (Pierce^TM^ DAB Substrate Kit; PI34002), for 4 minutes to avoid overstaining. Sections were air dried and dehydrated in an ascending series of ethanol/water solutions before being mounted with Paraclear (Polyscience; 22463) and a toluene solution. Images were taken using a Nikon Eclipse Ti-2 microscope with a standard light camera (Digital Sight 10). Quantification of DAB intensity was performed using ImageJ software. Briefly, images underwent color deconvolution using the ‘H DAB’ vector to isolate DAB staining. Background signals, measured from blank regions on each slide, were subtracted to enhance the accuracy of DAB staining quantification.

### Immunofluorescence

Dissected spinal cord slices (25 μm thickness, transverse plane and 20 μm thickness, sagittal plane) were washed in 1x PBS and permeabilized in a 1.5% PBS-Triton solution (Millipore-Sigma, USA) for 20 minutes. Blocking was performed using a 0.1% PBS-Triton solution containing 10% normal donkey serum (Millipore-Sigma, USA). The slices were then incubated overnight at 4℃ with the following primary antibodies: 1:1000 Iba-1 (Abcam, ab178846), 1:1000 NeuN (Abcam, ab104224), 1:1000 Tuj-1 (Abcam, ab18207), 1:1000 SARM-1 (Abcam, ab226930), and 1:1000 Myelin basic protein (MBP) (Abcam, ab40390). After incubation, tissue slices were stained with secondary antibodies (Alexa 488, Alexa 555, Alexa 594, and Alexa 647, Abcam, USA) at a 1:1000 dilution and mounted on coverslips using antifade mounting medium with DAPI (Vector, H2000). Images were captured using a Nikon Eclipse Ti-2 AX confocal microscope. Imaging areas within the target regions were randomly selected, and signal intensity was calculated and averaged from at least three slices per animal. Fluorescence intensity quantification was performed using ImageJ software.

### Locomotor Function

The recovery of locomotion deficits was evaluated using the Basso Mouse Scale (BMS), which was developed to assess open field locomotion deficits in spinal cord-injured mice. This scoring system is widely used as an indicator of recovery in mouse models of SCI. Observations were done on days 0, 1, 7, 14, 21, and 28 post-injury. Mice were observed in an open field for 5 minutes after they had gently adapted to the field. Left and right hindlimbs were assessed separately, and final scores were averaged.

### Mechanical hyperreflexia

Mechanical hyperreflexia was evaluated using calibrated von Frey filaments to quantify paw withdrawal thresholds in response to a known mechanical stimulus. Behavioral indices of neuropathic pain were measured before surgery as a baseline and repeated at 7, 14, 21, and 28 days after SCI. Briefly, animals were placed on an elevated metal mesh platform, covered by a transparent plastic container, and allowed to acclimate for 15 min before the assessment. A monofilament was pressed perpendicularly against the plantar surface of the hindlimb until bent, beginning with the 2.0 gf monofilaments and ranging from the 0.4 gf to 6.0 gf monofilaments. Each hindlimb was tested three times, and a two-thirds withdrawal threshold was calculated for each animal and reported as the average of both hind paws at each time point.

### Data Analysis and Statistics

The sample sizes were based on a two-sided power analysis performed in G*Power software on the experimental groups to allow us to detect a 10-unit difference in the most variable measurement at α=0.05 and β=0.8 based on the typical variance we have in our data. Statistical analysis was performed using Prism 10. Multiple comparisons were carried out by ordinary One-Way ANOVA or Brown-Forsythe and Welch ANOVA tests. The student’s t-test was used when comparing only two groups, where appropriate. p < 0.05 was considered statistically significant, and the results are expressed as the mean ± SEM.

## Results

### The expression and activity of ALDH2 in wild-type and ALDH2*2 mice with or without Alda-1 treatment in control and after SCI

To investigate the functional consequence of ALDH2 deficiency, we utilized an ALDH2*2 transgenic (TG) mouse model that carries a human ALDH2 inactivating mutation (E487K). Our initial experiments assessed ALDH2 protein expression and enzymatic activity in ALDH2*2 and wild-type (WT) mouse tissue after SCI and treatment with Alda-1, an ALDH2 activator, to evaluate the feasibility and gain necessary background information on ALDH2 for our study design. Immunoblotting and densitometric analysis (Fig. 2A) revealed that ALDH2 protein expression levels in the spinal cord were similar between ALDH2*2 homozygous and WT mice before and at three days post-trauma (*p* = 0.378, F = 8.848). We then conducted enzymatic activity assays to examine differences in ALDH2 efficiency and the activation effects of Alda-1 in spinal cord and liver tissues following SCI. ALDH2 enzymatic activity (Fig. 2B, C) was significantly higher in WT mice compared to ALDH2*2 mutants in uninjured groups, both in spinal cord (315.10 ± 23.01%, *p* = 0.009) and liver tissues (512.24 ± 72.92%, *p* = 0.109). Treatment with Alda-1 (10 mg/kg, twice daily) significantly enhanced ALDH2 enzymatic activity in the spinal cord three days post-SCI for ALDH2*2 (217.52 ± 17.4%, *p* = 0.0022) and WT (243.71 ± 17.17%, *p* = 0.0053) mice compared to untreated injured controls. Additionally, Alda-1 significantly increased ALDH2 activity in the liver tissue of ALDH2*2 mice by 311% (*p* < 0.001), but no such effect was observed in WT mice. These findings highlight a pronounced enzymatic deficiency in ALDH2*2 transgenic mice and demonstrate that Alda-1 effectively enhances ALDH2 activity following SCI in both genotypes.

**Figure 2.**
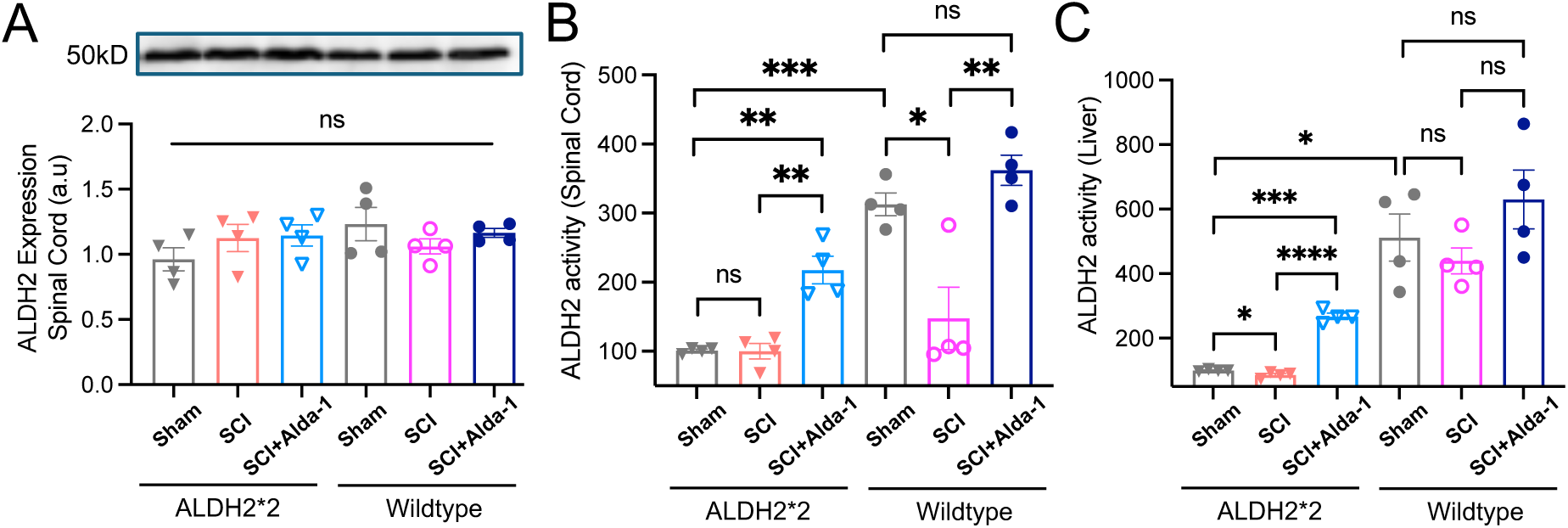
ALDH2 expression and activity in ALDH2*2 and wild-type mice following SCI with or without Alda-1. Mice were divided into sham, SCI, and “SCI + Alda-1” groups, with Alda-1 (10 mg/kg) administered intraperitoneally twice daily for 3 days post-SCI (6 total doses). (A) Representative Western blot images showing ALDH2 protein (∼ 50 kDa) expression across groups. ALDH2 expression levels were normalized by Coomassie brilliant blue total protein staining (Supplementary Fig. 1) and were comparable between ALDH2*2 mutants and wild-type sham, SCI, and SCI with Alda-1 treatment mice. No significant differences were observed between groups (*p* = 0.3783). (B, C) ALDH2 enzymatic activity, measured as Vmax (milli-units/min) at 5 min, in spinal cord and liver tissue lysates in the presence or absence of Alda-1. ALDH2 activity was significantly reduced in ALDH2*2 mutant mice compared to wild-type mice in the sham group in both tissues. Alda-1 treatment among the SCI groups significantly enhanced ALDH2 activity in both genotypes in the spinal cord, but only in the liver in the ALDH2*2 group, with greater effects observed in mutants. Data are presented as mean ± SEM (n = 3–4 per group). Statistical analysis was performed using one-way ANOVA followed by Tukey’s multiple comparisons test. *p < 0.05, **p < 0.01, ***p < 0.001, ****p < 0.0001.

### ALDH2*2 mutant mice exhibit exaggerated acrolein elevation in the spinal cord, mitigated by Alda-1 treatment over 28 days post-injury

The reactive aldehyde acrolein is a well-recognized pathological factor in SCI^11,12,46^. Acrolein levels, as indicated by DAB signal intensity, were quantified in the spinal cord over 28 days post-injury. In the SCI group, both WT and ALDH2*2 mice exhibited a significant increase in acrolein accumulation compared to their respective sham groups at 2 days (WT *p* < 0.0001, ALDH2*2 *p* < 0.0001), 7 days (WT *p* < 0.0001, ALDH2*2 *p* < 0.0001), and 28 days (WT *p* = 0.0043, ALDH2*2 *p* = 0.0017) post-injury (Fig. 3B, top # symbols). Two-way ANOVA analysis revealed that an ALDH2 deficiency exacerbated acrolein expression in the spinal cord compared to WT levels at 2 days (*p* = 0.0078, F = 8.974) and 7 days (*p* = 0.0469, F = 4.637) post-injury (Fig. 3C). These findings suggest that insufficient endogenous ALDH2 activity results in an impaired ability to clear reactive aldehydes, leading to heightened susceptibility to acrolein accumulation in the acute phase of SCI.

**Figure 3.**
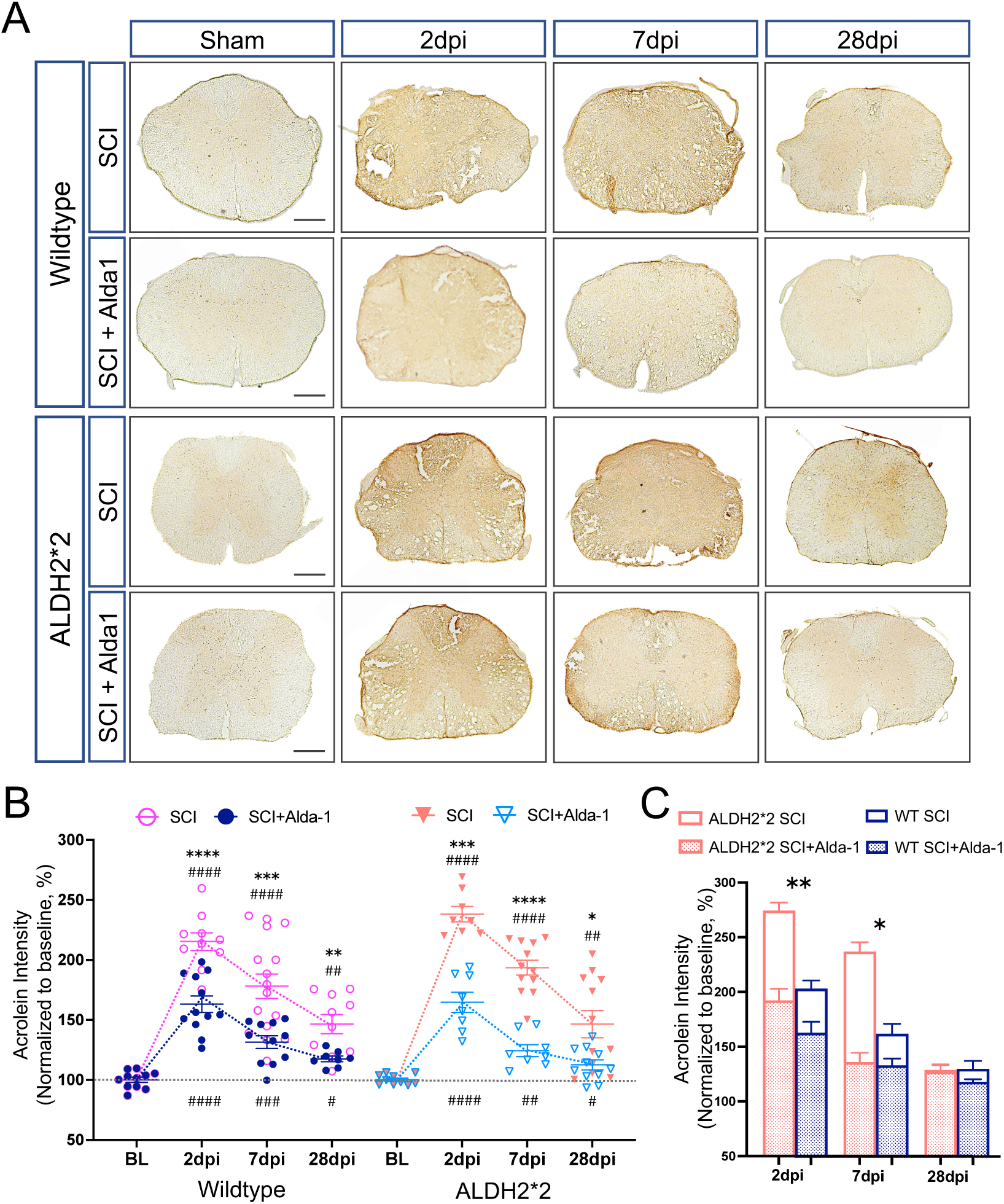
ALDH2*2 exacerbates acrolein accumulation in the spinal cord, which is alleviated by Alda-1 treatment. Mice were divided into sham, SCI, and “SCI + Alda-1” groups, with Alda-1 (10 mg/kg) administered intraperitoneally twice daily for the first 14 days post-SCI, beginning at 30 min post injury. (A) Representative transverse spinal cord images and (B) densitometric quantification of acrolein DAB staining demonstrate significantly elevated and sustained acrolein levels in the spinal cord of SCI groups compared to sham animals in both wild-type (WT) and ALDH2*2 mutant mice from 2 to 28 days post-injury. Alda-1 treatment significantly reduced acrolein accumulation in both genotypes across all time points. (C) ALDH2*2 mutant mice exhibited significantly greater acrolein accumulation at 2 and 7 days post-SCI compared to WT mice, indicating increased vulnerability due to ALDH2 deficiency. Note that no statistical difference was observed between WT and ALDH2*2 mice in the presence of Alda-1 across all time points. Data are presented as mean ± SEM (n = 8–16 per group). Statistical analysis was performed using Welch’s t-test, Brown-Forsythe Welch ANOVA, and Two-Way ANOVA. Significance was denoted as follows: *p < 0.05, **p < 0.01, ***p < 0.001, ***p < 0.0001. For (B), the symbol * represents significance between the SCI and SCI+Alda-1 groups, while # represents significance between either the SCI or SCI+Alda-1 and the sham (baseline) groups. For (C), the symbol * represents the significance for acrolein build-up (vulnerability) between the WT and ALDH2*2 mice after SCI.

Alda-1, an isozyme-specific activator of ALDH2, was administered intraperitoneally at a dose of 10 mg/kg twice daily for up to 14 days post-SCI in the treatment groups. Alda-1 treatment significantly reduced acrolein levels at 2 days (WT *p* < 0.0001, ALDH2*2 *p* < 0.0001), 7 days (WT *p* = 0.0005, ALDH2*2 *p* < 0.0001), and 28 days (WT *p* = 0.0055, ALDH2*2 *p* = 0.0133) post-injury (Fig. 3B, * symbols) in the spinal cord when compared to the untreated groups. However, acrolein accumulation persisted at higher levels than in control mice even after treatment at 2 days (WT *p* < 0.0001, ALDH2*2 *p* < 0.0001), 7 days (WT *p* = 0.0001, ALDH2*2 *p* = 0.0021), and 28 days (WT *p* = 0.03, ALDH2*2 *p* = 0.05) post-injury (Fig. 3B, bottom # symbols). These results indicate that acrolein elevation is prolonged after SCI, highlighting the need for continuous and effective strategies to control its toxic burden. Collectively, these findings indicate that the pharmacological enhancement of ALDH2 activity effectively mitigates the post-SCI acrolein surge, underscoring the critical role of ALDH2 in regulating toxic aldehyde accumulation following spinal cord injury.

### ALDH2 deficiency intensifies inflammation in the spinal cord, which is ameliorated by Alda-1 treatment at 7 days post-SCI

To assess immune cell activation, we stained sagittal spinal cord sections for active microglia/macrophages using the Iba-1 marker. The Iba-1-positive area was quantified and normalized to the total spinal cord area (%) (Fig. 4A, left; Fig. 4B). Post-injury, ALDH2*2 mice exhibited a trend of larger and more diffuse distribution of Iba-1-positive cells compared to WT mice (*p* = 0.077), suggesting a potentially more extensive and widespread inflammatory response and infiltration in the ALDH2*2 group. Alda-1 treatment effectively reduced both the area of immune activation and the number of activated microglia/macrophages (Fig. 4A, right; Fig. 4C) at the lesion epicenter in ALDH2*2 (*p* = 0.0039) and WT (*p* = 0.0162) mice at 7 days post-injury.

**Figure 4.**
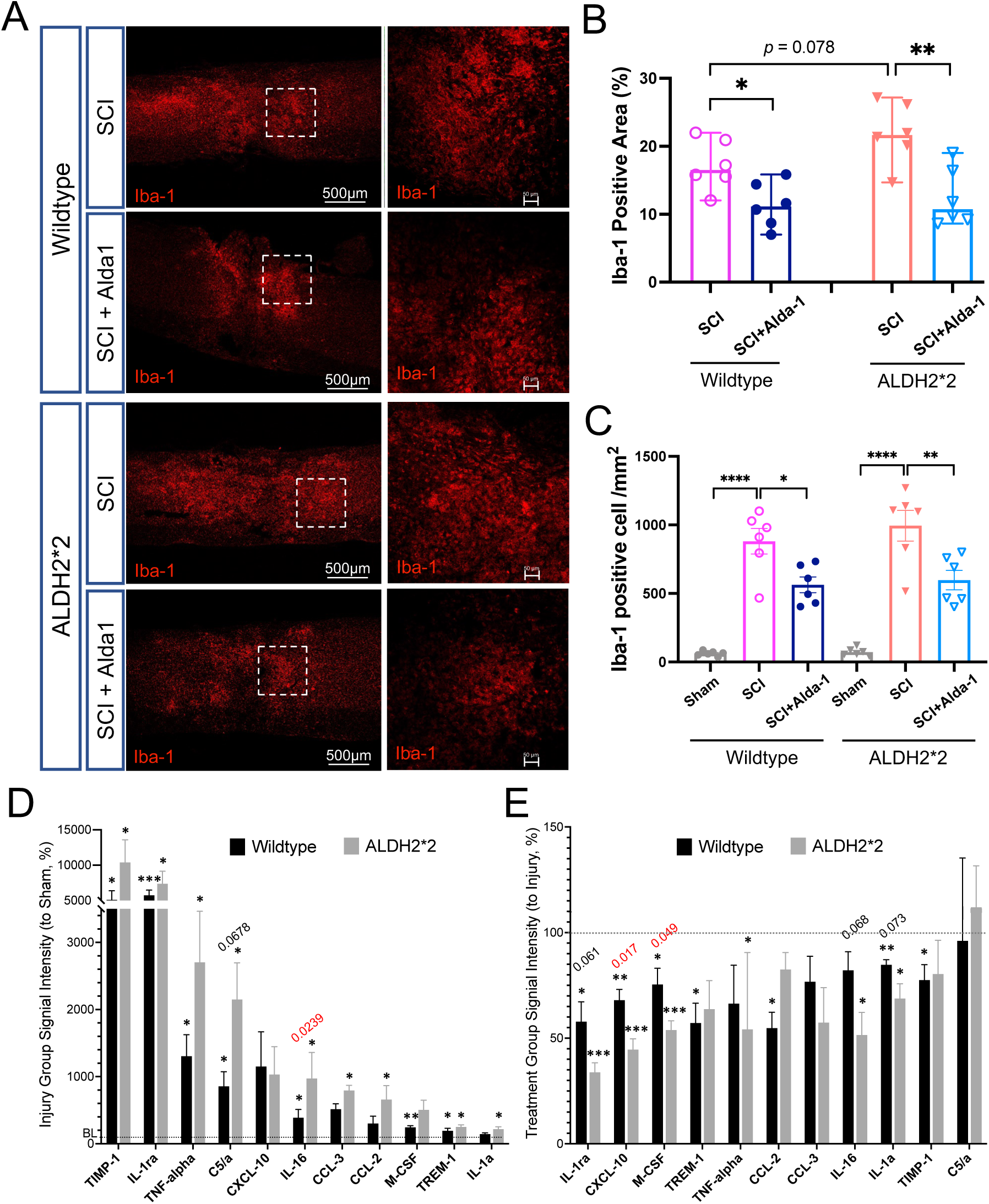
ALDH2 deficiency aggravates inflammation in the spinal cord, which is mitigated by Alda-1 treatment. (A) Representative images illustrate increased expression and activation of reactive microglia/macrophages, visualized using anti-Iba-1 (red), in the spinal cord post-SCI. Scale bars = 500 µm (low magnification) and 50 µm (high magnification). (B) Quantitative analysis of the Iba-1 positive area (normalized to the total spinal cord area, %) and (C) Iba-1-positive cell count revealed severe immune activation and widespread inflammation after SCI, while ALDH2*2 mice showed a trend toward exacerbated effects compared to WT mice (*p* = 0.078). Alda-1 treatment significantly reduced immune activation in both ALDH2*2 (*p* = 0.004) and WT (*p* = 0.016) mice after SCI. (D) Cytokine expression analysis showed a pronounced cytokine storm (normalized to sham, %) in the spinal cord at 7 days post-injury in both WT and ALDH2*2 mice, with ALDH2*2 mice exhibiting a tendency of more severe responses. (E) Alda-1 treatment effectively reduced levels of pro-inflammatory cytokines (normalized to untreated SCI group, %), particularly those associated with immune cell recruitment, with a greater impact observed in CXCL-10 and M-CSF in ALDH2*2 mice. All data are presented as mean ± SEM (n = 6 per group for immunofluorescence assays; n = 4 per group for cytokine assays). Statistical analyses included Brown-Forsythe Welch ANOVA with Tukey’s multiple comparisons test for group differences and two-way ANOVA for cytokine changes between genotypes. *p < 0.05, **p < 0.01, ***p < 0.001, ****p < 0.0001. For (D), symbol * represents the significance of the comparison between the SCI and sham groups in either WT or ALDH2*2 animals. For (E), symbol * represents the significance of the comparison between the SCI and treatment groups in WT or ALDH2*2 animals. Numbers of the p values are included when comparisons between the WT and ALDH2*2 groups are significant or nearly significant.

SCI triggers a massive and uncontrolled release of pro-inflammatory cytokines and other immune signaling molecules, amplifying the initial trauma^47,48^. Analysis of inflammatory cytokine expression in the spinal cord post-SCI revealed a pronounced cytokine storm at 7 days after injury in both WT and ALDH2*2 mice (Fig. 4D). Notably, the elevations of Complement Component 5a (C5a) (*p* = 0.0678) and Interleukin-16 (IL-16) (*p* = 0.0239), both critical for immune cell recruitment, were intensified in ALDH2*2 compared to WT mice. Treatment with Alda-1 markedly reduced systemic inflammatory signals in the spinal cord in both genotypes (Fig. 4E). Interestingly, Alda-1 treatment demonstrated a trend of greater reduction in the levels of Interleukin-1 alpha (IL-1α) (*p* = 0.073), Interleukin-Receptor Agonist (IL-1Ra) (*p* = 0.061), IL-16 (*p* = 0.068), C-X-C Motif Chemokine Ligand 10 (CXCL10) (*p* = 0.017), and Macrophage Colony-Stimulating Factor (M-CSF) (*p* = 0.049) in ALDH2*2 compared to WT mice. These findings suggest that a therapy to enhance aldehyde scavenging may reduce inflammatory responses after SCI by effectively modulating macrophage activity and mitigating Blood-Spinal Cord Barrier (BSCB) disruption.

### Alda-1 treatment protects against tissue loss post-SCI

Minimizing lesion size and promoting controlled scar formation are essential for facilitating axonal regrowth and neural plasticity, both of which are critical for the restoration of motor and sensory functions following SCI^49,50^. To investigate the pathological consequences of excessive acrolein accumulation after injury, we analyzed lesion histopathology in ALDH2*2 and WT mice. At 7 days post-injury, ALDH2-deficient mice exhibited a significantly larger lesion size, as indicated by the Tuj-1-negative area, compared to WT mice (*p* = 0.0206). Treatment with Alda-1 significantly reduced lesion size in ALDH2*2 mice (*p* = 0.0243) at the same time point (Fig. 5A, B). However, Alda-1-treated ALDH2-deficient mice revealed only a trend toward smaller lesion sizes at 28 days post-injury (*p* = 0.0694). Notably, Alda-1 treatment demonstrated greater efficacy in reducing gliosis in ALDH2*2 mice than in WT mice, particularly during the early phase following injury. This enhanced responsiveness may be attributed to the more pronounced aldehyde-induced pathology in ALDH2-deficient mice, rendering them more sensitive to therapeutic intervention targeting aldehyde detoxification.

**Figure 5.**
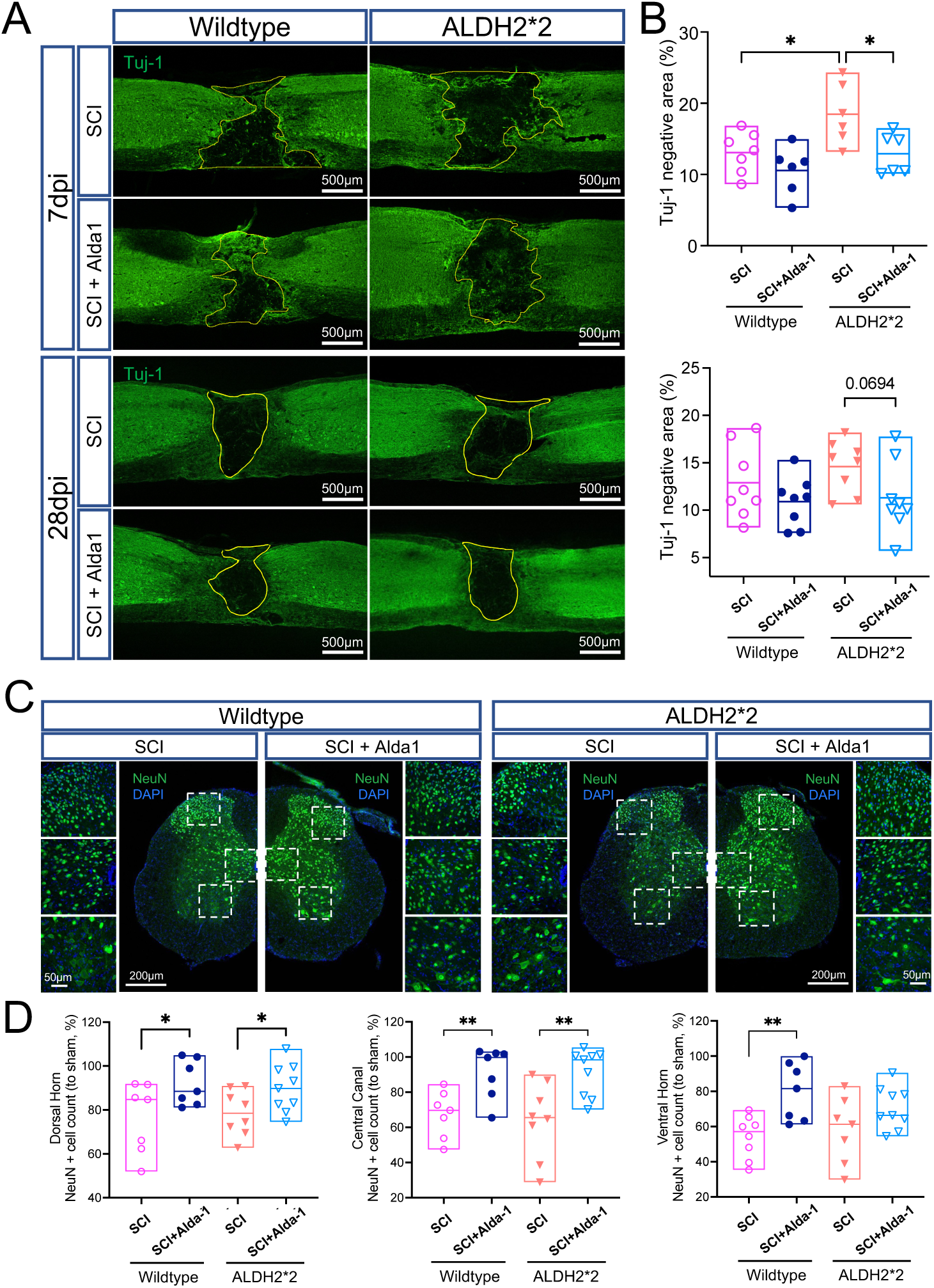
Alda-1 treatment protects against tissue loss following SCI. (A) ALDH2*2 mice show increased lesion size (compared to WT), while Alda-1 treatment reduces scar formation after SCI. Representative images of Tuj-1-negative lesion areas in the spinal cord of ALDH2*2 and WT mice at 7 and 28 days post-SCI, with lesion boundaries outlined in yellow. Scale bar = 500 µm. (B) Quantitative analysis of the Tuj-1-negative area (normalized to the total spinal cord area, %) revealed significantly larger lesion sizes in ALDH2*2 mutants compared to WT mice at 7 days post-SCI (top), while no difference was found at 28 days post-SCI (bottom). Alda-1 treatment in ALDH2*2 mice significantly reduced lesion size at 7 days while showing a strong tendency of reduction at 28 days. (C) Alda-1 treatment promotes neuronal survival in the spinal cord at 28 days post-SCI. Representative images of neurons labeled with anti-NeuN (green) in spinal cord sections slightly below the injury epicenter. Scale bar = 200 µm. Higher magnification views depict the dorsal horn, central canal, and ventral horn regions of the spinal cord. DAPI (blue) was used for nuclear counterstaining. Scale bar for higher magnification images = 50 µm. (D) Quantitative analysis of NeuN-positive cell counts (normalized to sham baseline) demonstrated that Alda-1 treatment, an acrolein scavenging strategy, significantly mitigated neuronal loss in multiple spinal cord regions in both ALDH2*2 and WT mice at 28 days post-SCI. Data are presented as mean ± SEM (n = 6–9 per group). Statistical analysis was performed using one-way ANOVA followed by Tukey’s multiple comparisons test. *p < 0.05, **p < 0.01.

Preserving neuronal viability and enhancing neuroprotection are critical strategies for maintaining spinal cord function and improving recovery outcomes after SCI^51^. As anticipated, neuronal survival—measured by NeuN-positive cell counts in the dorsal horn, central canal, and ventral horn—was markedly reduced in the SCI groups of both ALDH2*2 and WT mice (Fig. 5C, D). While no significant differences in neuronal survival were observed between the genotypes in the injured groups, Alda-1 treatment significantly mitigated neuronal cell death at 28 days post-injury in the dorsal horn (WT *p* = 0.0466, ALDH2*2 *p* = 0.0463) and central canal (WT *p* = 0.0077, ALDH2*2 *p* = 0.0059) for both WT and ALDH2*2 mice, but only for WT mice in the ventral horn (*p* = 0.0039, ALDH2*2 *p* = 0.0119).

### ALDH2 deficiency exacerbates demyelination in the spinal cord, whereas Alda-1 treatment preserves myelin integrity at the chronic stage post-SCI

The lipid-rich structure of the myelin sheath, combined with the oxidative environment following SCI, renders it highly susceptible to lipid peroxidation, which contributes to secondary injury mechanisms and exacerbates functional deficits^52,53^. Demyelination was assessed through immunohistochemical staining of myelin basic protein (MBP) in the spinal cord at 28 days post-injury. A significant reduction in MBP expression was observed in the ALDH2*2 SCI group, particularly in the lateral column area, compared to the WT group (Fig. 6B) (*p* = 0.0109). Alda-1 treatment effectively mitigated demyelination and preserved myelin integrity in the dorsal column area in both WT (*p* = 0.0198) and ALDH2*2 SCI groups (*p* = 0.0014), as evidenced by the retention of the characteristic circular morphology of myelin sheaths at high magnification (Fig. 6A, third row). These results highlight the critical role of acrolein overload in demyelination after SCI and illustrate the protective effects of Alda-1 in maintaining myelin integrity at 28 days post-injury.

**Figure 6.**
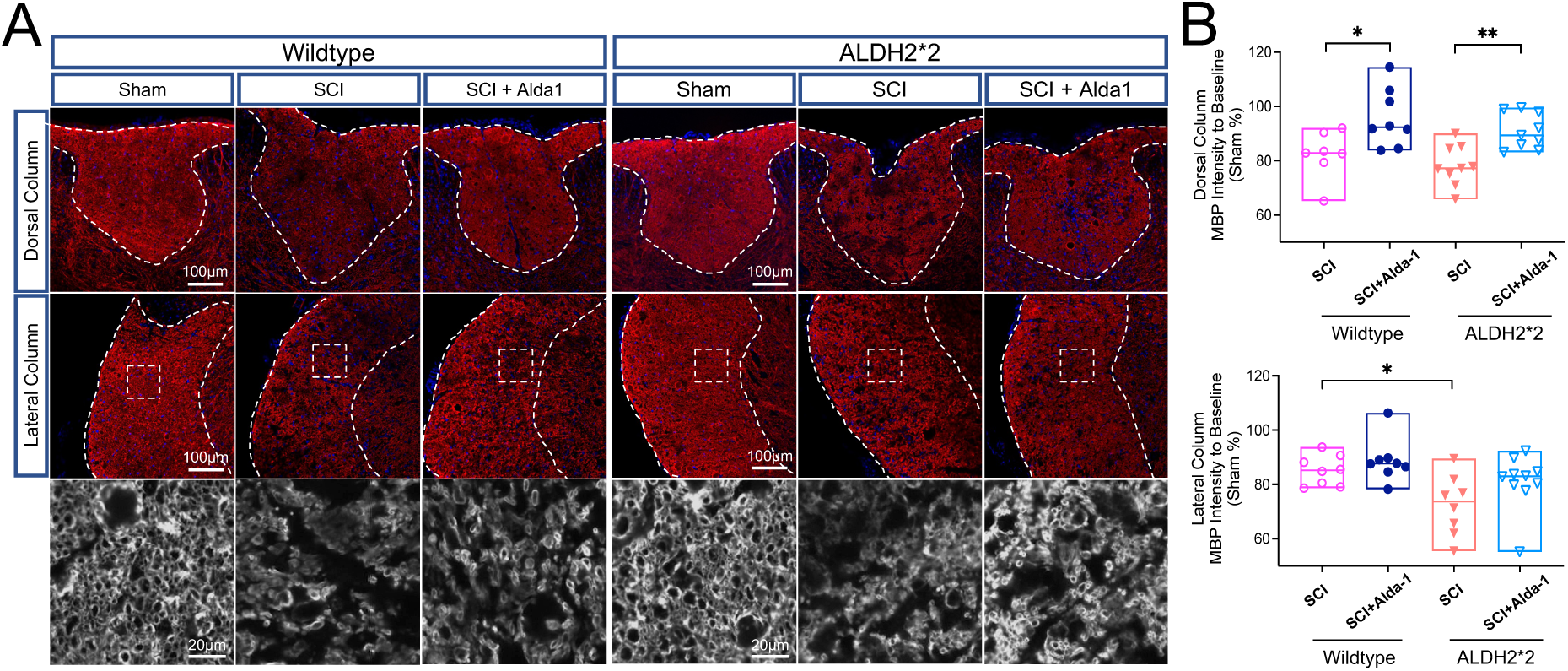
ALDH2*2 exacerbates demyelination, which is mitigated by Alda-1 treatment after SCI. (A) Representative images of myelin basic protein (MBP, red) and DAPI (blue) immunohistochemical staining in the spinal cord at 28 days post-injury. White matter regions, including the dorsal column and lateral column, are outlined in white (Scale bar = 100 µm). Higher magnification views (3^rd^ row) highlight morphological changes in axonal myelin sheaths within the lateral column. Scale bar = 20 µm. (B) Quantitative analysis of MBP signal intensity (normalized to sham baseline) revealed exacerbated demyelination in the lateral column of ALDH2*2 mice compared to WT mice (bottom). Alda-1 treatment significantly reduced myelin loss and axonal degeneration in the dorsal column regions of both WT and ALDH2*2 mice at 28 days post-injury (top and bottom). Data are presented as mean ± SEM (n = 7–9 per group). Statistical analysis was performed using one-way ANOVA followed by Tukey’s multiple comparisons test. *p < 0.05, **p < 0.01.

### Alda-1 treatment relieved locomotor and sensory impairments after SCI

The Basso Mouse Scale (BMS) locomotor rating scale was used to assess functional recovery and locomotor behavior in chronic SCI studies. Animals that suffered from SCI initially exhibited flaccid paralysis and scored 0-1 points immediately after the injury, with subsequent modest time-dependent recovery over 28 days (Fig. 7A, left). Notably, mice with an ALDH2 deficiency had lower average motor scores compared to WT mice at all timepoints post-injury, although no statistical significances were achieved (Day 7, *p* = 0.1, Day 14, *p* = 0.16). The ALDH2*2 mice that received Alda-1 treatment showed statistically significant improvements in lower-limb motor function from 7 to 28 days (p <0.05, 0.001, 0.05, 0.05 respectively) compared to the injury-only mice, while WT mice with treatment showed a significant recovery at the later stage of 21– and 28-days post-injury (p < 0.05 respectively). This data indicates a potentially more rapid and stronger compensation effect on motor functional recovery from the aldehyde-scavenging strategy in ALDH2-deficient individuals.

**Figure 7.**
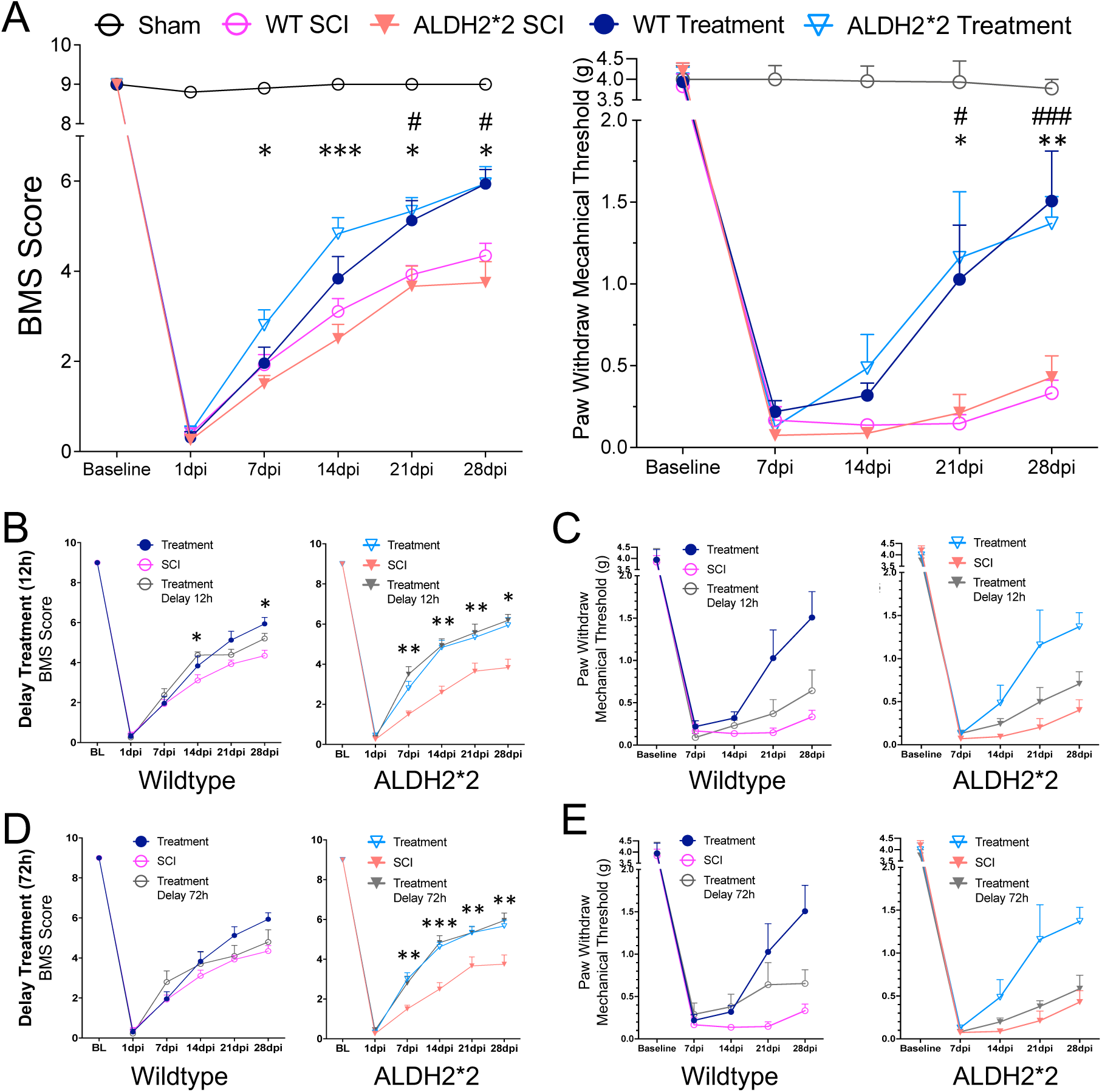
Enhancing ALDH2 activity alleviates locomotor and sensory impairments, even with delayed treatment. Time course analysis of SCI-induced motor and sensory dysfunctions with or without Alda-1 treatment. (A, left) Schematic of lower limb locomotor function assessment using the Basso Mouse Scale (BMS). Alda-1 treatment, beginning at 30 min post injury, progressively enhanced the recovery of lower limb motor function, showing more pronounced effects in ALDH2-deficient mice from day 7 to day 28 and in WT mice at later stages. (A, right) Schematic of sensory function assessment using the von Frey filament (VF) test for pain threshold evaluation. Mice with SCI exhibited severe and persistent mechanical hyperalgesia, which was significantly mitigated by Alda-1 treatment in both ALDH2*2 and WT mice at 21 and 28 days post-SCI. (B, C) BMS scores and VF test results in mice receiving their delayed first Alda-1 dose at 12 hours post-injury showed improved motor and sensory outcomes compared to untreated controls. (D, E) BMS scores and VF test results in mice receiving their delayed first Alda-1 dose at 72 hours post-injury also demonstrated significant motor functional recovery. Data are presented as mean ± SEM. For (A), n = 10–15 per group, while n = 7–8 per group for delayed treatment (B-E). Statistical significance was determined using Brown-Forsythe Welch ANOVA, followed by Tukey’s multiple comparisons test. For (A), the symbol * indicates comparisons between injury and treatment groups in ALDH2*2 mice, while # indicates comparisons in WT mice. * indicates comparisons between the injury and delayed treatment groups for (B-E). *p < 0.05, **p < 0.01, ***p < 0.001.

Neuropathic pain is a common symptom following SCI in both human patients and animal models^54,55^. Mechanical allodynia was assessed using the von Frey filament test (VF) on the hind limbs before and at 7-day intervals after SCI (Fig. 7A, right). Before sham and SCI surgery, all animals’ average paw withdrawal thresholds reached the cut-off of 4g. Mice that underwent spinal cord injury showed a significant drop in mechanical thresholds starting at day 7, indicating hypersensitivity to noxious stimuli. In contrast, both ALDH2*2 and WT animals receiving Alda-1 treatment exhibited significant recovery from hyperreflexia at 21 (WT *p* = 0.0298, ALDH2*2 *p* = 0.0497) and 28 days (WT *p* = 0.0046, ALDH2*2 *p* = 0.0004) post-injury. Therefore, Alda-1 appears to effectively alleviate both motor deficits and neuropathic pain following SCI.

To evaluate the effectiveness of the ALDH2 rescue strategy under more clinically relevant conditions, we included two delayed treatment groups, where the first dose of Alda-1 was administered at either 12 or 72 hours post-injury. Notably, Alda-1 treatment significantly improved lower limb motor function in ALDH2*2 mice at 7, 14, 21, and 28 days post-injury, when treatment initiation was delayed to 12 (*p* = 0.0016, *p* = 0.0037, *p* = 0.0052, *p* = 0.0131) or 72 hours post-injury (*p* = 0.001, *p* = 0.0008, *p* = 0.0043, *p* = 0.004) (Fig. 7B, D), highlighting a relatively extended therapeutic window for the therapeutic enhancement of neuronal functional recovery. In contrast, WT mice exhibited motor recovery only at 14 (*p* = 0.0256) and 28 days (*p* = 0.0387) post-injury when treatment was initiated within 12 hours, but not at 72 hours post-SCI. The von Frey (VF) test revealed a trend toward recovery from mechanical allodynia in both ALDH2*2 and WT mice across both delayed treatment groups, although these differences did not reach statistical significance (Fig. 7C, E). These findings suggest that, even beyond the traditionally ideal treatment window, an aldehyde-scavenging therapy remains effective in promoting functional recovery after SCI, particularly in motor recovery and to a greater extent in ALDH2*2 mice.

## Discussion

In this study, we utilized an ALDH2*2 transgenic mouse model with an inactive form of the enzyme to investigate the impact of a compromised endogenous aldehyde-scavenging system on pathological changes and neurofunctional recovery after SCI. Our findings revealed that ALDH2 deficiency resulted in pronounced acrolein accumulation, which was associated with heightened inflammatory responses, tissue loss, and demyelination in the spinal cord compared to WT animals. Treatment with Alda-1, an ALDH2 activator, enhanced ALDH2 activity and significantly reduced acrolein levels in the spinal cord of both ALDH2-deficient and WT mice from 2 to 28 days post-SCI. This reduction in acrolein levels was accompanied by diminished microglial activation, suppression of the cytokine storm, reduced neuronal death, and improved preservation of myelin integrity. These effects translated to significant improvements in motor and sensory functions up to 28 days post-injury. In particular, the neuroprotective effects of therapeutically enhancing ALDH2 activity were often more pronounced in the ALDH2*2 mice. To our knowledge, this is the first study to comprehensively evaluate the critical role of ALDH2 activity in SCI by both up– and down-manipulating its function, as well as examining different therapeutic time windows beyond the traditional acute-stage period for broader application.

Under physiological conditions, acrolein is primarily metabolized and detoxified by mitochondrial aldehyde dehydrogenase 2 (ALDH2) through enzymatic mechanisms^37,56^. However, excessive acrolein production post-injury can overwhelm or inactivate this detoxification system. Acrolein forms stable adducts with nucleophilic residues in the active site of ALDH2, impairing its catalytic activity and perpetuating a vicious cycle of toxicity^57,58^. In addition to obstructing the active site of ALDH2, studies have suggested that the reduced aldehyde clearance observed in the spinal cord after trauma or in cardiac tissue following ethanol consumption may also result from direct damage to the enzyme^29,59^. In contrast, other studies have reported that ALDH2 expression levels remain unchanged across different injury models^30,60^. In our study, statistical analysis revealed no significant difference in ALDH2 levels pre– and post-injury across all groups (Fig. 2A). These findings suggest that the inefficiency of acrolein scavenging after SCI is likely driven primarily by a reduction in ALDH2 catalytic activity rather than a decrease in enzyme quantity.

Our study represents the first to measure ALDH2 activity in the spinal cord under ALDH2-deficient genetic conditions, revealing a two-thirds reduction in enzymatic activity compared to WT animals (Fig. 2B). Alda-1 restores ALDH2 function by binding at the entrance of the catalytic tunnel, facilitating substrate catalysis^61^. Previous studies have reported that Alda-1 enhances ALDH2 catalytic activity by 11-fold in the cardiac tissue of ALDH2*2 homozygous mice^37^. In our study, Alda-1 significantly increased ALDH2 activity by more than twofold in the spinal cord and threefold in the liver after three days of treatment in ALDH2*2 mice following SCI (Fig. 2B, C). These findings align with studies conducted on hepatic and brain tissues and suggest that the pharmacological efficacy of Alda-1 may vary across different tissue types^62,63^.

Despite the significant accumulation of acrolein that occurs following SCI, we observed that Alda-1, an ALDH2 enhancer, can significantly reduce levels of acrolein at all timepoints post-injury (Fig. 3A, B). Furthermore, the crippled ALDH2 in our ALDH2*2 mice appeared to exaggerate acrolein elevations at the early stages following SCI (Fig. 3C). Importantly, boosting ALDH2 activity led to a reduction in acrolein levels in both ALDH2*2 and WT animals, with a significantly greater effect observed in ALDH2*2 mice after SCI. These findings not only underscore the essential role of ALDH2 in acrolein metabolism but also position it as a compelling therapeutic target—distinct from conventional aldehyde scavengers—for mitigating acrolein-associated pathologies. Moreover, given that ALDH2*2 is among the most prevalent human genetic polymorphisms, these results suggest that individuals carrying this variant may experience more severe SCI outcomes.

Among secondary injury mechanisms, excessive activation of the systemic immune response and neuroinflammation are key contributors to injury progression^3^. Reactive oxygen species (ROS) and lipid peroxidation products, such as acrolein, act as signaling molecules that activate microglia, astrocytes, and infiltrating macrophages, promoting the release of pro-inflammatory cytokines and further amplifying inflammation^7,47^. In our study, increased macrophage/microglia activation (Iba1) correlated with elevated acrolein levels at 7 days post-SCI, reinforcing the link between neuroinflammation and impaired aldehyde detoxification (Fig. 4A, B, C). Furthermore, ALDH2*2 mice exhibited exacerbated levels of IL-1α, IL-1Ra, IL-16, CCL2, CCL3, CXCL10, and M-CSF in the spinal cord after SCI (Fig. 4D). These cytokines and chemokines are known contributors to SCI pathology through monocytes, macrophages recruitment and regulating glial cells, and their therapeutic neutralization has been shown to reduce inflammation and neuropathic pain in rodent models^64,65^. Although it remains unclear whether reduced macrophage infiltration and cytokine expression are direct consequences of acrolein removal or secondary effects of attenuated inflammatory activation, our findings do suggest that aldehyde scavenging mitigates neuroinflammation by limiting macrophage recruitment after SCI. The observed reduction in macrophage/microglia activation and cytokine levels following Alda-1 treatment indicates that acrolein plays a key role in sustaining the inflammatory response. Given that lipid peroxidation-derived aldehydes, including acrolein, can serve as damage-associated molecular patterns (DAMPs) that trigger immune activation^66,67^, it is plausible that their removal disrupts the positive feedback loop of inflammation. This suggests that acrolein scavenging may act upstream of macrophage recruitment by preventing the initial inflammatory cascade, thereby reducing both immune cell infiltration and the subsequent release of pro-inflammatory mediators. Crucially, acrolein scavenging through Alda-1 treatment significantly reduced macrophage/microglia activation and cytokine storms (Fig. 4B, C, E). This finding not only supports our proposed mechanism by which anti-acrolein strategies can mitigate inflammation— by inhibiting macrophage recruitment—but also underscores the potential synergistic benefits of simultaneously targeting oxidative stress and inflammatory mediators in SCI treatment.

In addition to neuroinflammation, we assessed tissue loss through multiple approaches. Acrolein-induced neurodegeneration and neuronal loss are well-documented in various neurodegenerative diseases, including animal models of SCI^12,22,68–70^. To address this, we evaluated neuronal loss in key regions of the spinal cord: the ventral horn, central canal (intermediate laminar), and dorsal horn (Fig. 5C). At 28 days post-SCI, significant neuronal loss was observed in all three regions in both ALDH2*2 and wild-type (WT) animals, as indicated by a marked reduction in NeuN-positive cell counts (Fig. 5D). However, no significant difference in neuronal loss was observed between ALDH2*2 and WT mice. Interestingly, although neurons at the dorsal horn region are reported to be more vulnerable to excitotoxicity and inflammation after SCI^71,72^, we observed the most dramatic neuronal loss (∼40%) in the ventral horn compared to the sham group, followed by the dorsal horn (∼20%) and central canal (∼15%). This heightened vulnerability of motor neurons in the ventral horn may be attributed to their higher metabolic rate and energy demand, rendering them more susceptible to oxidative stress and mitochondrial dysfunction^73,74^. Furthermore, MBP expression levels were significantly reduced in the SCI groups, particularly in the lateral column of ALDH2*2 mice. Alda-1 treatment exhibited a selective neuroprotective effect, rescuing MBP expression primarily in the dorsal column region (Fig. 6A, B). Collectively, these findings demonstrate that acrolein scavenging via Alda-1 treatment mitigates SCI-induced neurotoxicity, providing neuroprotection and promoting neurorepair. These effects are strongly correlated with enhanced ALDH2 activity and reduced acrolein levels.

Locomotor deficits, resulting from the combined effects of lesion-induced damage, neurodegeneration, neuroinflammation, and oxidative stress, are a significant consequence of SCI^1^. Consequently, the restoration of locomotor function remains a primary goal of preclinical studies^75^. In our study, a gradual and significant functional recovery was observed in both ALDH2*2 and WT mice within 28 days post-injury, supporting the therapeutic potential of Alda-1 (Fig. 7A). This recovery is likely associated with acrolein removal and anti-inflammatory effects, as discussed earlier. However, while both ALDH2*2 and WT SCI-only mice exhibited significantly lower BMS scores compared to their respective treatment groups, ALDH2*2 mice showed only an insignificant trend toward worse motor dysfunction relative to WT mice. To gain more precise insights into motor function recovery, additional evaluations such as gait analysis and electromyography (EMG) could be incorporated in future studies.

Although the neuronal survival in the ventral horn was more robust in WT mice treated with Alda-1 (Fig. 5D), motor improvements in ALDH2*2 mice receiving Alda-1 treatment were more rapid and superior compared to WT mice. We hypothesize that several factors may contribute to this phenomenon. First, studies using animals carrying the ALDH2*2 mutation have shown more efficient therapeutic effects in alleviating nociceptive responses and atrial fibrosis in other injury models compared to WT animals^76,77^. We speculate that neurons surviving after injury in ALDH2*2 mice may be more sensitive to enzymatic boosting by Alda-1, leading to a stronger compensatory response. Second, while a balance between excitatory and inhibitory neurons is crucial for functional recovery, excitatory neurons are reported to play a more critical role in maintaining locomotor function after SCI^78^. Enhancing ALDH2 activity may preferentially preserve excitatory neuronal integrity and function, thereby contributing to the improved locomotor outcomes observed in these animals. This differential sensitivity between neuronal subtypes to oxidative stress and detoxification pathways may underlie the enhanced therapeutic response in ALDH2-deficient models. Third, a specific population of excitatory interneurons located in the intermediate laminae at the thoracic level, which project to the lumbar level, has been identified as a key component in lower limb locomotor recovery after SCI^79,80^. The significant preservation of neuronal survival in the intermediate laminae region with Alda-1 treatment may facilitate the restoration of locomotor function. Future studies focusing on neuronal subpopulations could provide further insights into these mechanisms.

Neuropathic pain resulting from SCI has profound and long-term consequences, often surpassing the impact of other functional disabilities and severely diminishing patients’ quality of life^54^. Although the mechanisms underlying neuropathic pain are multifactorial and complex, studies have demonstrated a correlation between acrolein levels and nociceptive behaviors^12,70^. In our study, Alda-1 treatment significantly reduced pain-like behaviors at 21 and 28 days post-injury when initiated 30 min after the injury for the first two weeks post SCI, as measured by VF-filament mechanical tests (Fig. 7B). This suggests that Alda-1-induced activation of ALDH2 and suppression of acrolein levels regulate nociception and mitigate sensory hypersensitivity. These effects may be mediated through the modulation of neuropathic pain-associated mediators, including the pro-algesic transient receptor potential ankyrin 1 (TRPA1) and IL-16, as observed in previous studies^81–83^.

Early intervention within 8-24 hours post-SCI is critical for improving outcomes by mitigating secondary injury^84^. Surgical decompression and other early interventions are most effective when administered within this timeframe^85^. However, many patients face barriers to receiving timely care, underscoring the need for improved emergency response systems and access to specialized trauma centers^86^. Several animal studies have suggested that even delayed intervention could still contribute to functional recovery. Nakashima et al. reported that injection of the high mobility group box-1 (HMGB1) protein, aimed at reducing sterile inflammation, improved neuronal function in SCI mice when administered within 6 hours post-injury^87^. Similarly, Arnold’s study demonstrated that anti-inflammatory treatments−initiated at 6 weeks post-injury−could still yield temporary improvements in locomotor function^88^. Our previous study also illustrated that administering an acrolein scavenger starting from 21 days post-injury still mitigated hyperreflexia in SCI rats during the treatment period^69^. To explore the clinical potential of boosting ALDH2 function under different, and perhaps more clinically relevant and realistic scenarios, we evaluated whether delayed treatment initiated at 12 or 72 hours post-injury could still alleviate motor and sensory impairments. Surprisingly, significant improvements were observed in both ALDH2*2 and WT SCI mice when the first dose of Alda-1 was delayed by 12 hours, and even by 72 hours in ALDH2*2 mice (Fig. 7B, D). These results not only support our hypothesis that neurons controlling motor function in ALDH2*2 mice are more sensitive to enzymatic boosting and exhibit a stronger compensatory response to treatment, but also highlight the potential of an extended treatment window for an endogenous acrolein scavenging therapy. Unfortunately, only a non-significant trend toward improved sensory recovery was observed in both ALDH2*2 and WT mice in the delayed treatment groups (Fig. 7C, E).

Our study has several limitations that warrant consideration. First, the experiments were conducted exclusively on male mice, reflecting the higher incidence of SCI in males (male-to-female ratio ∼2:1). However, sex has been considered a critical biological variable that influences neuropathy, recovery, and therapeutic efficacy in SCI^89,90^. Future studies should include female subjects to evaluate sex-specific responses to ALDH2 activation and Alda-1 treatment, ensuring broader clinical applicability and a more comprehensive understanding of its therapeutic potential. Second, while our findings provide valuable insights, additional assessments such as single-cell RNA sequencing or detailed gait analysis at the cellular and behavioral levels could further elucidate genotype-specific differences and offer deeper mechanistic insights into Alda-1’s neuroprotective effects. Third, DMSO was used as a solvent for Alda-1 due to its solubility requirements. Although vehicle injections were administered to sham animals to control for potential biases, DMSO is known to cause gastrointestinal distress and other adverse effects when administered intraperitoneally^91^. Future studies should explore alternative administration routes, such as oral delivery with a more biocompatible solvent, to improve tolerance and minimize potential side effects.

## Conclusion

In conclusion, this study provides a comprehensive investigation into the impact of the ALDH2*2 genotype on SCI pathogenesis using a novel transgenic mouse model. Our findings demonstrate that ALDH2 deficiency leads to significantly increased oxidative stress, inflammation, tissue loss, demyelination, and behavioral impairments in ALDH2*2 mice compared to WT mice after SCI. The use of the ALDH2 activator Alda-1 highlights the critical role of ALDH2 in acrolein detoxification and suggests its potential as a therapeutic agent for mitigating SCI-induced damage. Notably, enhancing endogenous acrolein-scavenging machinery may be particularly beneficial for individuals carrying the ALDH2*2 mutation, offering a promising avenue for translational applications in SCI treatment.

## Abbreviations

DAMPs: damage-associated molecular patterns
M-CSF: macrophage colony-stimulating factor
PhzA: phenelzine analog
PUFAs: polyunsaturated fatty acids
TG: transgenic
Tuj-1: Anti-Beta-Tubulin III
WT: wild-type
dpi: days post injury

## Acknowledgments

We thank Dr. Che-Hong Chen at Stanford University for providing the ALDH2 transgenic mouse line as a gift. We also thank the current and prior members of the Shi laboratory for critical discussions and day-to-day assistance with the experiments included in this manuscript.

## Author contributions

This study is based on an original idea of SS, SH, and RShi. SS, TD, and RS carried out animal and biochemical studies. AA, TD, and ZZ critically examined the animal and biochemical studies. SS, AA, and RShi wrote and edited the manuscript. RShi contributed to the conceptualization, funding, overall supervision, supported review development, and overall editing. All authors approved of the final manuscript.

## Conflicts of interest

Riyi Shi is a co-founder of Neuro Vigor, a company developing novel drug treatments and diagnostic approaches for neurodegenerative diseases and neurotrauma. The remaining authors declare that they have no conflict of interests.

## Funding sources

The study was supported by a grant from National Institute of Neurological Disorders and Stroke R21 (R21NS115094).

## Ethics approval and consent for participation

We affirm that this paper contains original data that have not been submitted elsewhere for publication and that all authors have read and approved the manuscript. We have striven to follow ARRIVE Essential 10 guidelines in all aspects of this study. Animal procedures were approved by the Institutional Animal Care and Use Committee (IACUC) of Purdue University (No. <u>1909001957</u>) on December, 2023. Human data or human tissue has not been used in this study.

## Availability of data and materials

Datasets analyzed during the current study are available from the corresponding author on reasonable request.

---

**Supplementary Figure 1.**
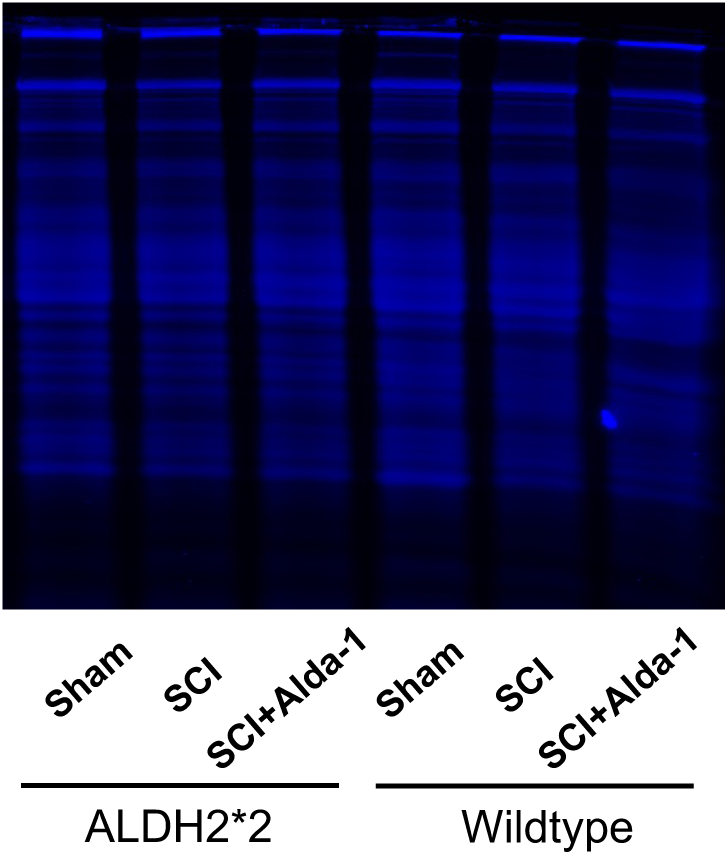
Coomassie Brilliant Blue staining of total protein as a loading control for spinal cord ALDH2 Western blot. ALDH2 expression was normalized to total protein (y-axis) in ALDH2*2 mutant and wild-type mice under sham, SCI, and SCI + Alda-1 treatment conditions (n = 4 per group).

